# DOT1L primarily acts as a transcriptional repressor in hematopoietic progenitor cells

**DOI:** 10.1101/2020.10.15.341255

**Authors:** Shaon Borosha, Anamika Ratri, Sami M. Housami, Shubham Rai, Subhra Ghosh, Carrie A. Malcom, V. Praveen Chakravarthi, Jay L. Vivian, Timothy A. Fields, M.A. Karim Rumi, Patrick E. Fields

## Abstract

DOT1L is essential for early hematopoiesis but the precise mechanisms remain largely unclear. The only known function of DOT1L is histone H3 lysine 79 (H3K79) methylation. We generated two mouse models; a *Dot1L*-knockout (*Dot1L*-KO), and another possessing a point mutation in its methyltransferase domain (*Dot1L*-MM) to determine the role of its catalytic activity during early hematopoiesis. We observed that *Dot1L*-KO embryos suffered from severe anemia, while *Dot1L*-MM embryos showed minimal to no anemia. However, *ex vivo* culture of *Dot1L*-MM hematopoietic progenitors (HPCs) exhibited defective development of myeloid and mixed progenitors. DOT1L is a well-recognized, cell-type specific epigenetic regulator of gene expression. To elucidate the mechanisms underlying such diverse hematopoietic properties of *Dot1L*-KO and *Dot1L*-MM HPCs, we examined their whole transcriptomes. Extensively self-renewing erythroblast (ESRE) cultures were established using yolk sac (YS) cells collected on embryonic day 10.5 (E10.5). *Dot1l-*KO and *Dot1l-*MM cells expanded significantly less than the wildtype cells and showed slower progression through the cell cycle. Total RNA extracted from the wildtype and *Dot1l-*mutant ESRE cells were subjected to RNA-seq analyses. We observed that the majority (~82%) of the differentially expressed genes (DEGs) were upregulated in both of the *Dot1L*-mutants, which suggests that DOT1L predominantly acts as a transcriptional repressor in HPCs. We also observed that about ~40% of the DEGs were unique to either of the mutant group, suggesting that DOT1L possesses both methyltransferase domain-dependent and -independent functions. We further analyzed Gene Ontology and signaling pathways relevant to the DEGs common to both mutant groups and those that were unique to either group. Among the common DEGs, we observed upregulation of CDK inhibitors, which explains the cell cycle arrest in both of the *Dot1L*-mutant progenitors.

## 1. INTRODUCTION

DOT1L histone methyltransferase (DOT1L) is essential regulator of vital tissue and organ development during embryonic life, including hematopoiesis^1^. We observed that loss of DOT1L in mice (*Dot1L*-KO) results in lethal anemia during mid-gestation^1^. DOT1L is the only known methyltransferase in eukaryotic cells to methylate lysine 79 of histone H3 (H3K79)^2^. We generated a mouse line carrying a point mutation (Asn241Ala) in mouse *Dot1L* gene (*Dot1L*-MM) that renders its catalytic domain inactive^3,4^. *Dot1L*-MM mice contained an intact DOT1L protein that lacked only the H3K79 methyltransferase activity^3^. The methyltransferase mutant, *Dot1L*-MM mice, were also embryonic lethal-died around mid-gestation^3^. The mice also displayed defects in embryonic hematopoiesis, including a decreased ability to form definitive myeloid, and oligopotent (mixed) blood progenitors in *ex vivo* cultures^3^. However, unlike the *Dot1L* knockout (*Dot1L-*KO)^1^, HPCs from the *Dot1L-*MM YS were able to produce erythroid colonies in numbers similar to the wildtype^3^.

Histone methylation is important for permissive or repressive chromatin conformation and can have a profound effect on regulation of gene expression^5^. DOT1L is responsible for the mono, di- and tri-methyl marks on lysine 79 of histone H3 (H3K79)^2^. These histone modifications as well as the DOT1L protein have been strongly associated with actively transcribed chromatin regions^6^. Thus, it has been suggested that DOT1L is involved in epigenetic regulation of transcriptional activation of genes in a tissue specific manner.

In this study, we examined the expression of DOT1L-regulated genes on embryonic day 10.5 (E10.5) yolk sac (YS) derived hematopoietic progenitor cells (HPCs). We observed that more than 82% of the differentially expressed genes (DEGs) in *Dot1L*-KO or *Dot1L*-MM HPCs cultured *ex vivo* were upregulated, which suggests that DOT1L primarily acts as a transcriptional repressor in HPCs.

## 2. METHOLODOLOGY

### 2.1. Dot1L mutant mouse models

The *Dot1L*-KO mice were generated and maintained as described previously^1^. To produce the *Dot1L*-MM mouse, we generated mutant mESC as described^3^. *Dot1L*-KO and *Dot1L-*MM heterozygous mice were maintained by continuous backcrossing to 129 stocks. Genotyping was performed on tail clips by using RED extract-N-Amp Tissue PCR Kit, Sigma-Aldrich as previously described^7,8^. All animal experiments were performed in accordance with the protocols approved by the University of Kansas Medical Center Animal Care and Use Committee.

### 2.2. Extensively Self-Renewing Erythroblasts (ESRE) assays

*Dot1L*-KO or *Dot1L*-MM heterozygous mutant males and females were set up for timed mating to collect the conceptuses on E10.5. Pregnant females were sacrificed, and uteri were dissected to separate embryos and YS. Embryos were treated with RED extract-N-Amp Tissue PCR reagents (Millipore Sigma, Saint-Louis, MO) to purify genomic DNA and perform the genotyping PCR^7,8^. Digested E10.5 YSs were washed in IMDM alone, resuspended in 0.5ml expansion media for ESRE according to England et al.,^9^, and plated into gelatin-coated 24 well plates. Expansion media consisted of StemPro34 supplemented with nutrient supplement (Gibco/BRL), 2 U/ml human recombinant EPO (University of Kansas Hospital Pharmacy), 100 ng/ml SCF (PeproTech), 10^−6^ M dexamethasone (Sigma), 40 ng/ml insulin-like growth factor-1 (PeproTech) and penicillin-streptomycin (Invitrogen). After 1 day of culture, the nonadherent cells were aspirated, spun down, resuspended in fresh ESRE media, and transferred to a new gelatin coated well. After 3 days in culture, RNA was extracted from wildtype, *Dot1L*-KO or *Dot1L*-MM HPCs.

### 2.3. Assessment of cell proliferation, cell cycle analyses and apoptosis assays

Single-cell suspensions from E10.5 YS were cultured in MethoCult™ GF M3434 (StemCell Technologies, Vancouver, BC, Canada) for 4 days. The mix of cytokines in this methylcellulose medium promotes definitive erythroid, myeloid, and mixed progenitor differentiation. Cells were collected on day 4 and stained with Annexin V to assess apoptosis. Some cells were fixed by adding cold 70% ethanol slowly to single cell suspensions, and then stained with propidium iodide^10^. Flow cytometry was performed by the use of a FACSCalibur (BD Biosciences, San Jose, CA)^11^. Analyses of the cytometric data were carried out using CellQuest Pro software (BD Biosciences)^12–14^.

### 2.4. Sample collection, library preparation and RNA-sequencing

RNA quality was assessed by a Bioanalyzer at the KUMC Genomics Core, and samples with RIN values over 9 were selected for RNA-sequencing library preparation. RNA samples were extracted from multiple, expanded YS cells obtained from embryos of the same genotype. These samples were pooled to prepare each RNA-seq library. Approximately 500 ng of total RNA was used to prepare a RNA-seq library using the True-Seq mRNA kit (Illumina, San Diego, CA) as described previously^15–17^. The quality of RNA-seq libraries was evaluated by Agilent Analysis at the KUMC Genomics Core and the sequencing was performed on an Illumina NovaSeq 6000 sequencer (KUMC Genomics Core).

### 2.5. RNA-seq data analyses

RNA-sequencing data were demultiplexed, trimmed, aligned, and analyzed using CLC Genomics Workbench 12.2 (Qiagen Bioinformatics, Germantown, MD) as described previously^15–17^. Through trimming, low-quality reads were removed, and good-quality reads were aligned with the *Mus musculus* genome (mm10) using default guidelines: (a) maximum number of allowable mismatches = 2, (b) minimum length and similarity fraction = 0.8, and (c) minimum number of hits per read = 10. Gene expression values were measured in transcripts per million (TPM). DEGs were identified that had an absolute fold change of TPM ≥ 2 and a false discovery rate (FDR) *p*-value of ≤0.05.

### 2.6. Gene Ontology (GO) and disease pathway analyses for the RNA-sequencing data

DEGs were subjected to Gene Ontology (GO) analysis (http://www.pantherdb.org) and categorized in biological, cellular and molecular function. DEGs in *Dot1L-*mutant HPCs were further analyzed by Ingenuity Pathway Analysis (IPA; Qiagen Bioinformatics, Germantown, MD) to build gene networks related to placental development. Functional analyses were performed towards understanding the biological pathways and functions altered in either of the *Dot1L-*mutant progenitor cells.

### 2.7. Validation of RNA-sequencing data

DEGs were validated by RT-qPCR. RT-qPCR validation included cDNA samples prepared with wildtype, *Dot1L*-MM and *Dot1L*-KO ESRE cell derived total RNAs. The genes were selected from the IPA analyses and MGI data that impacted the proliferation and differentiation of HPCs.

### 2.8. Statistical analysis

Each RNA-seq library or cDNA was prepared from pooled RNA samples extracted from at least 3 different ESRE cultures of the same genotype. Each group for RNA sequencing consisted of three independent libraries and the DEGs were identified by CLC Genomics workbench as described previously^15–17^. RT-qPCR validation included at least six cDNA samples prepared from wildtype, *Dot1L*-MM and *Dot1L*-KO ESRE cell total RNA. The experimental results are expressed as mean ± standard error (SE). The RT-qPCR results were analyzed by one-way ANOVA, and the significance of mean differences was determined by Duncan’s *post hoc* test, with *p* ≤ 0.05. All the statistical calculations were done using SPSS 22 (IBM, Armonk, NY).

## 3. RESULTS

### 3.1. Dot1L-KO and Dot1L-MM embryos exhibit distinct hematopoietic phenotypes

We observed that *Dot1L*-KO embryos develop slower than WT embryos and suffer from lethal anemia ^1^ (**Fig.1A-E**). *Dot1L*-KO embryos die between embryonic day 11.5 (E11.5) and E13.5. *Ex vivo* culture of HPCs from E10.5 *Dot1L*-KO YS showed that erythroid differentiation was severely affected compared to myeloid lineage ^1^. We generated another mouse model that carries a point mutation (Asn241Ala) in endogenous DOT1L, rendering the catalytic domain inactive^18^ (**Fig.1F-J**). Although the *Dot1L*-methyl mutant (*Dot1L-*MM) embryos also died at midgestation, we observed remarkable differences in the hematopoietic phenotype between the *Dot1L*-KO and *Dot1L*-MM mice^18^ (**Fig.1B-E** and **G-J**); in particular, erythropoiesis was minimally affected in *Dot1L*-MM YS and embryo, suggesting that hematopoietic activity of DOT1L may not be limited to its MT domain. However, ex vivo culture of YS cells exhibited that formation of myeloid and mixed colonies were dramatically reduced in either *Dot1L*-KO or *Dot1L*-MM^18^. Culture of *Dot1L*-KO HPCs showed decreased cell proliferation (**Fig. 2A**), accumulation of cells in G_0_/G_1_ stage (**Fig. 2B**), and a greater percentage of *Dot1L*-KO or *Dot1L*-MM HPCs in ESRE culture were Annexin V-positive ^1^ (**Fig. 2C**). In addition, Alkaline Comet assays showed a greater DNA damage in *Dot1L-*MM compared to WT cells and the DNA damage was still higher in *Dot1L-*KO ESREs compared to MM (**Fig. 2 D**).

**Fig.1.**
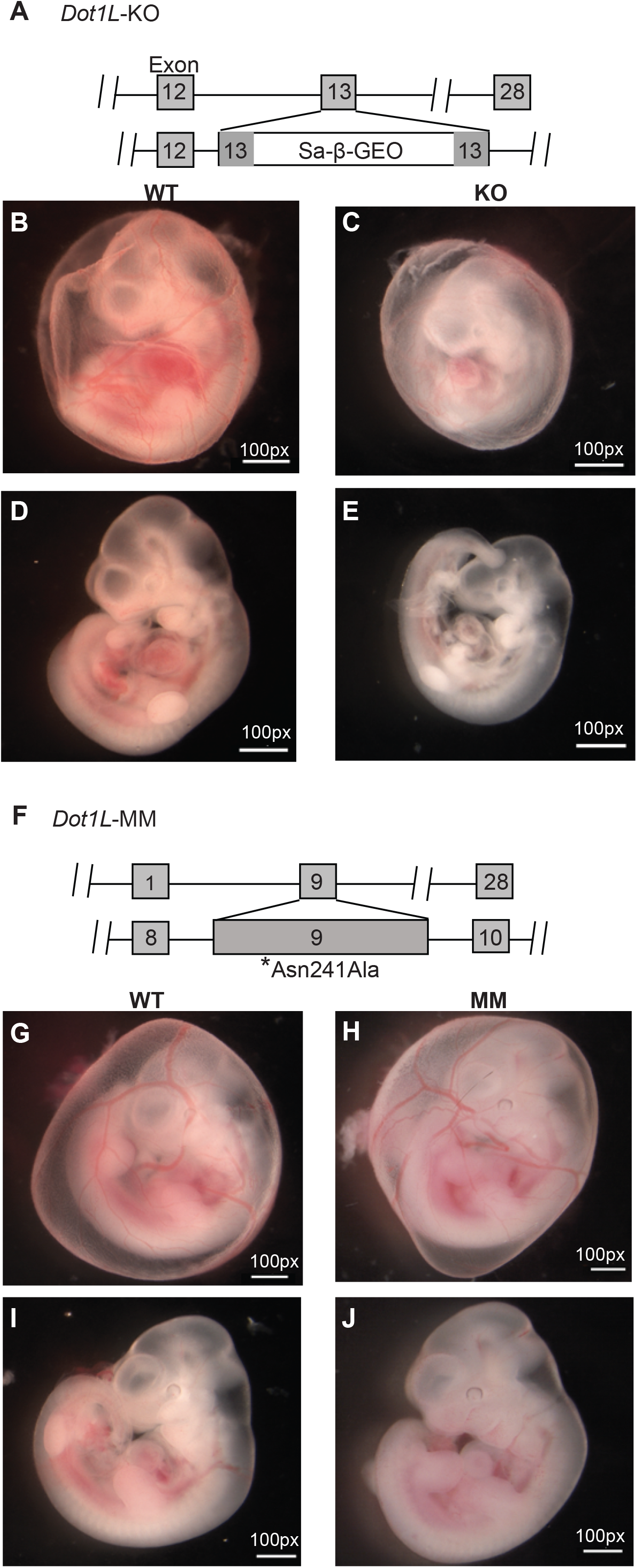
Differential phenotypes of *Dot1L*-KO and *Dot1L*-MM embryos. Schematic representation of the gene trap construct used to generate the mutant *Dot1L*-KO alleles (**A**). Representative images of E10.5 wildtype (WT) (**B, C**), *Dot1L*-KO (**D, E**), show that growth of *Dot1L*-KO embryos was slower than that of WT (**B-E**). **F**) Schematic diagram of mouse *Dot1L* gene targeted by CRISPR/Cas9 to introduce a point mutation (Asn241Ala) that eliminates its MT activity (*Dot1L*-MM). Representative images of E10.5 WT (**G,H**), *Dot1L*-MM (**I,J**), show that growth of *Dot1L*-MM embryos was comparable to that of WT. While *Dot1L*-MM embryos displayed embryonic lethality, they had little to no anemia, unlike KO (**B-J**). Vasculature of KO yolk sac has noticeably less blood than that of either the WT or MM YS (**B-J**).

**Fig. 2.**
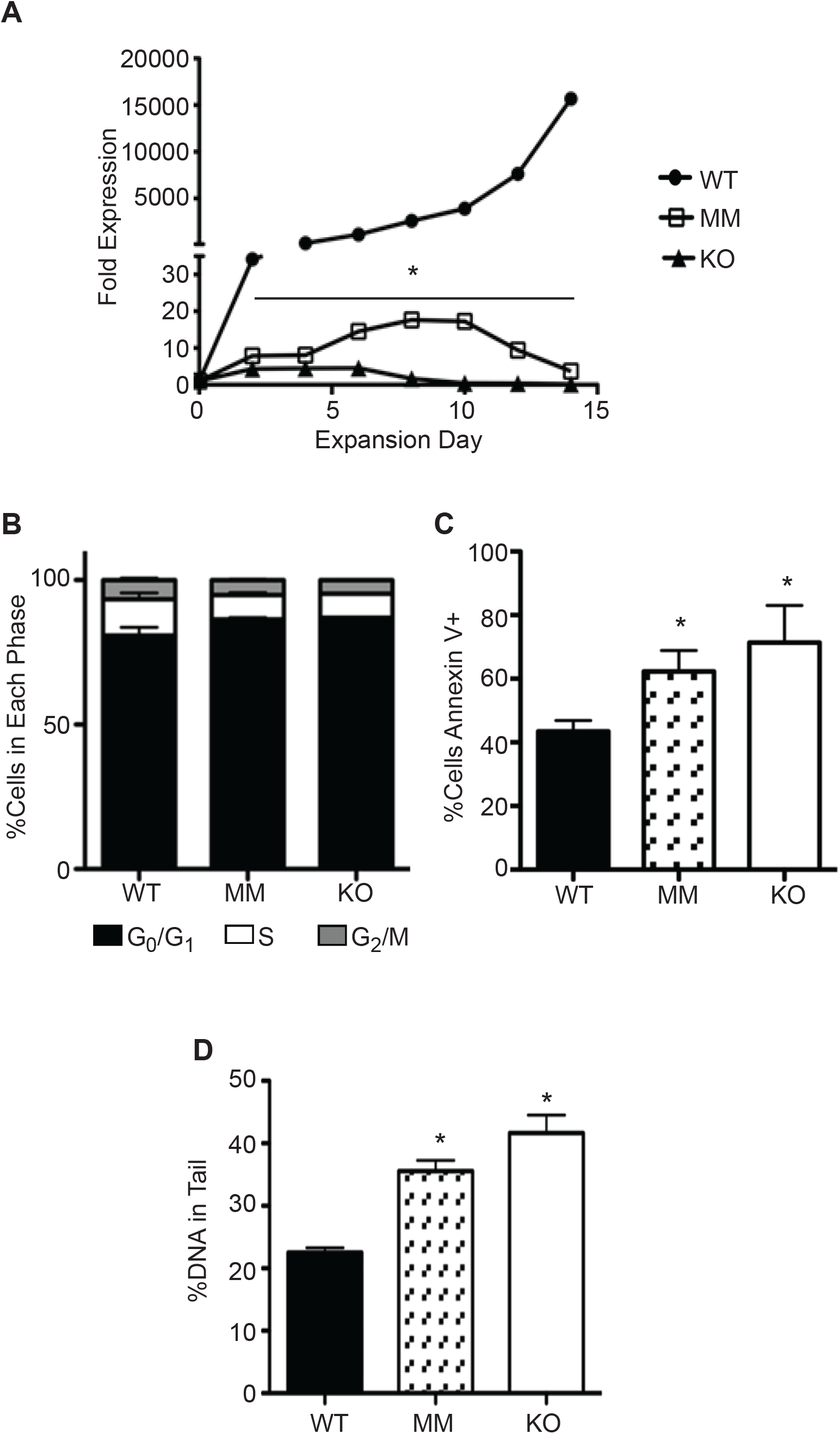
*Dot1L*-KO and *Dot1L*-MM ESREs display defective proliferation and survival. Cells isolated from E10.5 wildtype (WT), *Dot1L*-KO, and *Dot1L*-MM YSs were cultured in expansion medium for ESREs. (**A**) Cell number was counted via trypan blue exclusion every 2 days for 14 days, and fold expansion was calculated relative to day 3 after isolation. On day 6-7 of counting, cells were labeled for propidium iodide or Annexin V and analyzed via flow cytometry for cell cycle analysis (**B**) and apoptosis (**C**). Alkaline Comet assays were performed on day 6 ESREs to assess DNA damage. *Dot1L*-KO, *Dot1L*-MM and ESREs contained greater levels of DNA damages compared to WT (**D**).

### 3.2. DEGs in Dot1L-KO or Dot1L-MM ESRE cells

Transcriptome data-sets were generated by sequencing of mRNA purified from ESRE cultures using E10.5 wildtype, *Dot1L*-KO or *Dot1L*-MM YS cells. The raw data have been deposited to NCBI SRA under PRJNA666736. Analyzed data include the DEGs are shown in **Fig. 3(A-F)**. Of the total 25,749 reference genes in the mm10 genome, 16,806 genes were detected in wildtype and 16,678 in *Dot1L*-KO and 17,053 in *Dot1L*-MM ESRE cells. Analyses of the detected genes for level of gene expression revealed that ~40% had a very low abundance (<1 TPM), ~20% had low abundance (1-5 TPM), ~12% had modertate abundance (>5-10 TPM), ~25% had high abundance (>10-100 TPM), and only ~3% of the genes had a very high abundance (> 100 TPM). Among these genes, 2238 were differentially expressed (absolute fold change ≥2, *p*-value ≤ 0.05) in *Dot1L*-KO, with 358 downregulated and 1698 upregulated. In contrast, 2752 genes were differentially expressed (absolute fold change ≥2, *p*-value ≤ 0.05) in *Dot1L*-MM cells, with 328 downregulated and 1649 upregulated. The DEGs were evident in the hierarchical clustering (**Fig. 3A, B**) and Volcano plots (**Fig. 3C, D**), which demonstrate that most of the DEGs were upregulated in *Dot1L*-KO or *Dot1L*-MM HPCs (**Fig. 3E, F**).

**Fig. 3.**
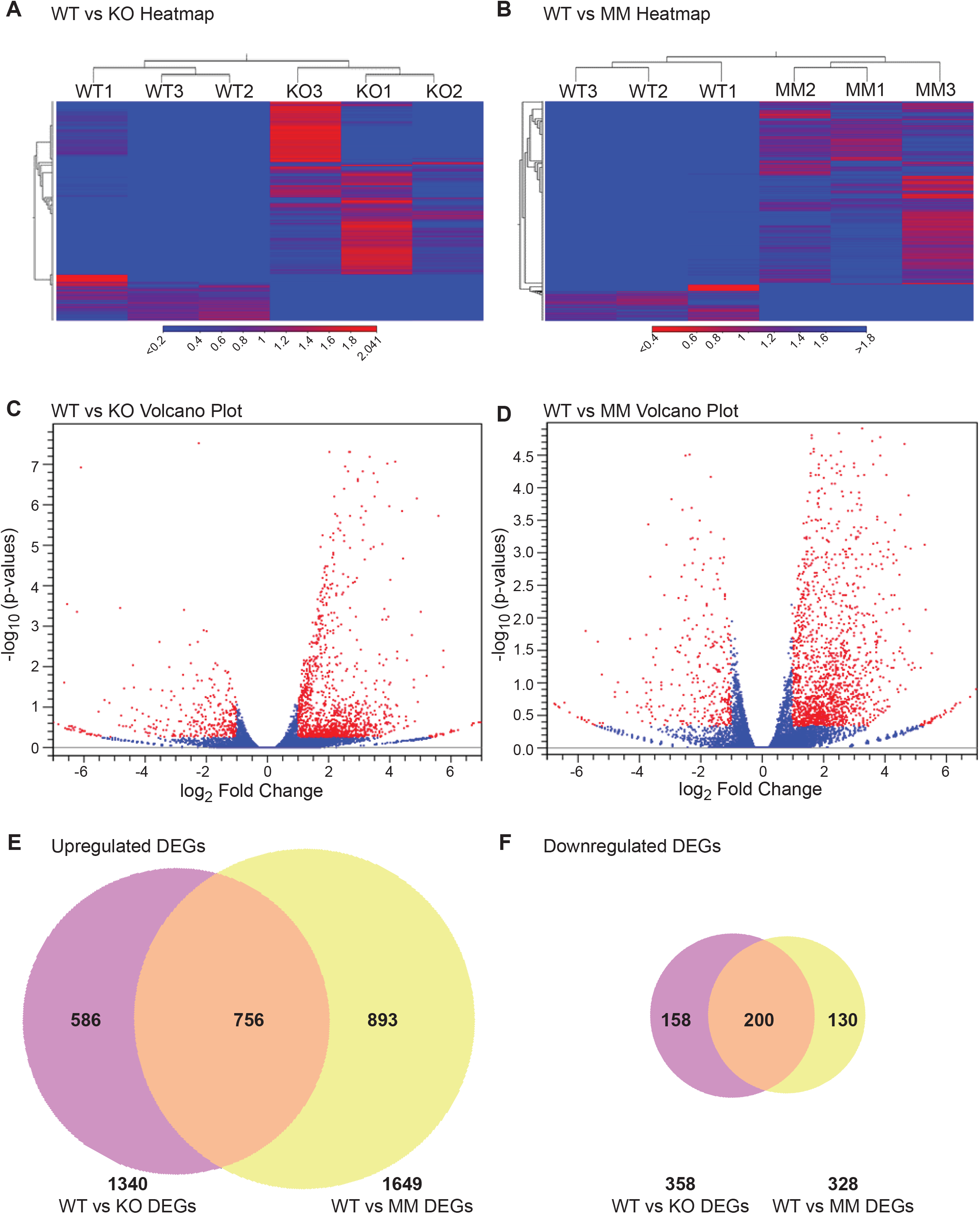
DEGs *Dot1L*-KO and *Dot1L*-MM ESREs. Analyses of HPC transcriptomes were performed in wildtype (WT), *Dot1L*-KO, and *Dot1L*-MM ESRE on the 3rd day of *ex vivo* culture. RNA-seq libraries were prepared using 500ng of total RNAs extracted from pooled ESRE cells and sequencing was performed on an Illumina NovaSeq 6000 platform. RNA-seq data were analyzed using the CLC Genomics Workbench. All expressed genes were distributed according to TPM values. Hierarchical clustering was performed on differentially expressed genes (absolute fold change ≥ 2, *p* value ≤ 0.05) in WT versus *Dot1L*-KO groups (**A**), and WT versus *Dot1L*-MM groups (**B**). The DEGS in the *Dot1L*-KO, and *Dot1L*-MM ESRE cells are presented in a Volcano plots (**C, D**respectively), with red dots showing the differentially expressed genes (n=3/genotype). Finally, Venn diagrams showing that majority of the DEGs (~82%) in either *Dot1L*-KO or *Dot1L*-MM are upregulated (**E, F**). Venn diagrams also showed that about 35-40% of the DEGs were unique to either *Dot1L*-mutant groups (**E, F**).

### 3.4. Gene Ontology (GO) analyses of the DEGs

GO analysis classified the DEGs into three categories: Biological process (**Fig. 4A, B**), Molecular function (**Fig. 4C, D**) and Cellular component (**Fig. 4E, F**). GO analysis revealed that the majority of the genes in the biological process group were involved in biological process, cellular processes, or cell signaling (**Fig. 4A, B**). The genes in molecular function were involved in binding, protein-protein interactions, catalytic activity, and molecular and transcriptional regulation (**Fig. 4C, D**). The genes in cellular component were predominantly involved in cell parts, membranes, organelles, and protein-protein complexes (**Fig. 4E, F**).

**Fig. 4.**
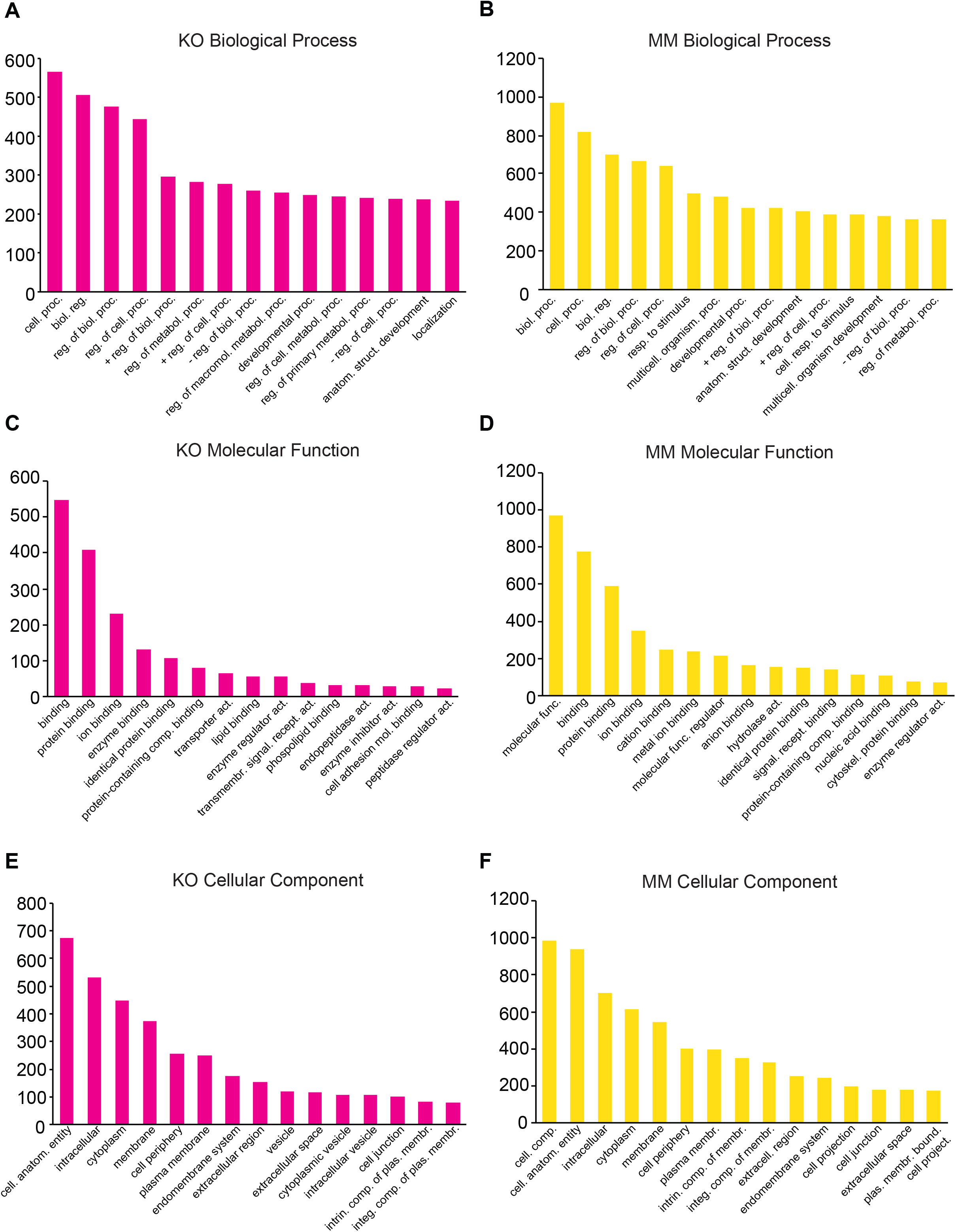
Classification of differentially expressed genes based on Gene Ontology. The differentially expressed genes in *Dot1L*-KO and *Dot1L*-MM ESREs were subjected to Panther Classification Analysis (http://pantherdb.org). Differentially expressed genes were classified based on Biological process (**A**), Molecular function (**B**) and Cellular components (**C**).

### 3.5. Ingenuity Pathway Analysis (IPA) of the DEGs

IPA of the DEGs in *Dot1L*-KO or *Dot1L*-MM HPCs in ESRE culture identified upregulation of genes related to regulation of hematopoiesis. Among the hematopoietic pathways involved, we were particularly interested in proliferation and differentiation of hematopoietic progenitor cells (**Fig. 5A-D**). The DEGs included upregulation of CDK inhibitors and downregulation of *Flt1*, *Flt3*, *Hoxa9* and *Mpl*. Interestingly, among the 165 known genes involved in differentiation of HPCs, only 9 were affected in *Dot1L*-KO cells and 21 in *Dot1L*-MM ESRE cells (**Fig. 5 E, F**).

**Fig. 5.**
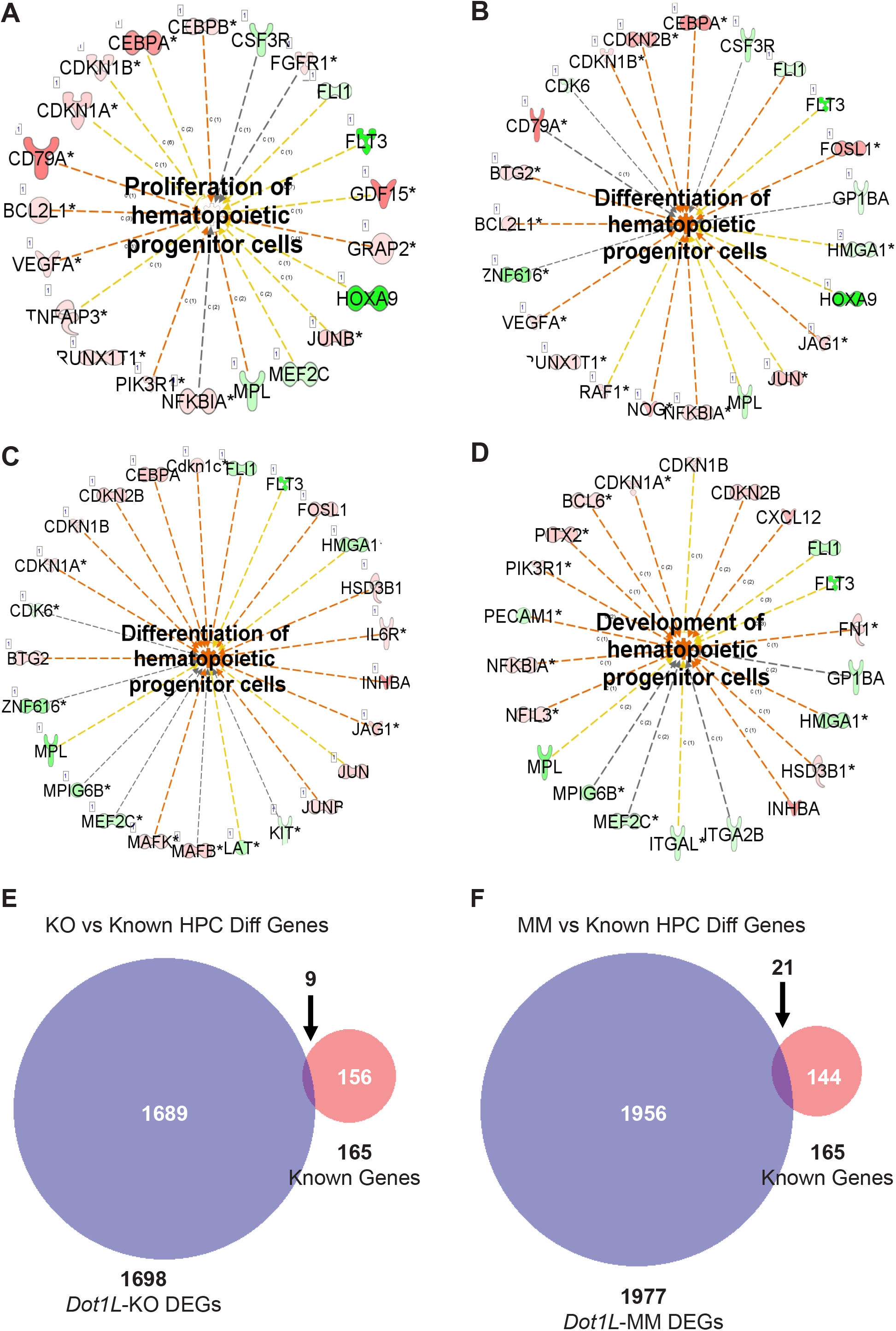
DEGs involved in proliferation and differentiation of HPCs. DEGs from RNA-seq data representing wildtype (WT) versus *Dot1L*-KO (**A, B**), and WT versus *Dot1L*-MM (**C, D**) were subjected to IPA analysis and detected many genes crucial for proliferation, and differentiation HPCs. In addition, we curated a group of genes (n=165) from MGI database (www.informatics.jax.or) that are involved in differentiation of HPCs and compared those with the DEGs identified in our RNA-seq data. Most of DEGs in *Dot1L*-KO or *Dot1L*-MM ESRE cells were found to be novel.

### 3.6 RT-qPCR analyses validated DEGs involved in HPC proliferation and differentiation

Differentially expressed genes that were identified to be involved in proliferation and differentiation of HPCs were validated by RT-qPCR analyses. We observed that while genes involved in proliferation of HPCs were significantly downregulated in both *Dot1L*-KO and *Dot1L*-MM ESRE cells (**Fig. 6 A-F**), those involved in induction of differentiation were markedly upregulated (**Fig. 6 G-I**). Marked increase in CDK inhibitors (**H, I**) can explain the accumulation of cells in G_0_/G_1_ phase and increased proportion of the *Dot1L*-mutant ESRE cells undergoing apoptosis.

**Fig. 6.**
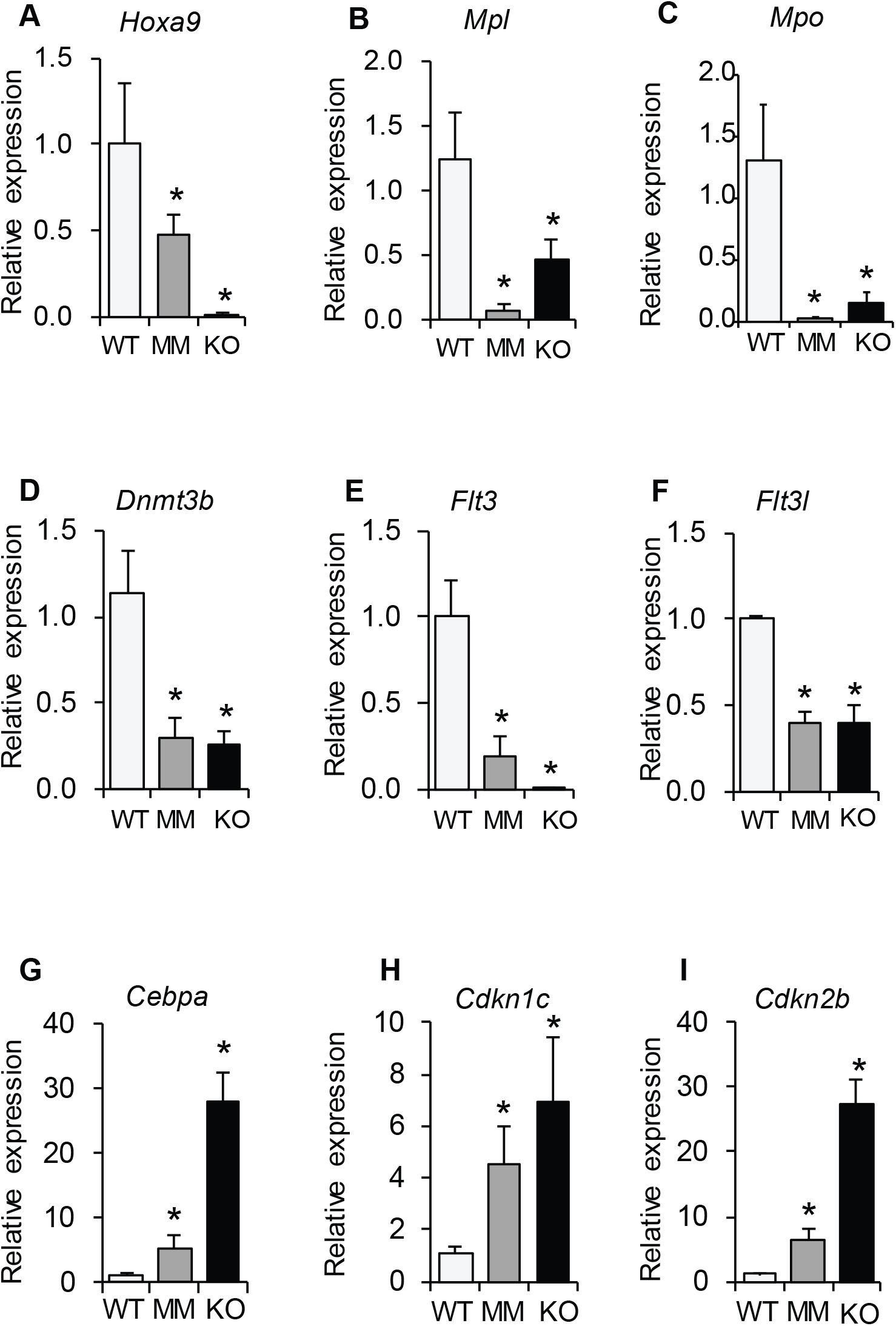
RT-qPCR validation of selected DEGs involved in proliferation and differentiation of HPCs. Differentially expressed genes that were identified to be involved in proliferation and differentiation of HPCs were further validated by RT-qPCR analyses. We observed that while genes involved in proliferation of HPCs were significantly downregulated in both *Dot1L*-MM and *Dot1L*-MM ESRE cells (**A-F**). In contrast, those involved in induction of HPC differentiation were markedly upregulated (**G-I**). Marked increase in CDK inhibitors (**H, I**) can explain the increased cell number in G_0_/G_1_ phase and increased percentage of the *Dot1L*-mutant ESRE cells undergoing apoptosis.

## 4. DISCUSSION

DOT1L is expressed at high levels in mouse HPCs (**Supplemental Fig.1**), suggesting a potential role for this chromatin organizer and transcriptional regulator in early blood development. This study analyzed the DEGs in both *Dot1l-*KO and *Dot1L-*MM mouse HPCs derived from E10.5 YS to examine early blood development. RNA-seq datasets were used to identify DOT1L-regulated genes in mouse HPCs and understand their potential role in early blood development.

We previously reported that hematopoietic transcription factor (TF) *Gata2* was significantly reduced in *Dot1L-*KO HPCs, whereas *Pu.1*, an erythropoiesis inhibiting TF was upregulated^1^. We also observed that KIT-positive HPCs from *Dot1L*-KO YS expressed low levels of *Trpc6*^19^. Our RNA-seq data recapitulated the previous observations regarding expression of *Gata2, Pu.1 and Trpc6* (PRJNA666736).

DOTL1L is responsible for methylation of H3K79^2^. Histone methylation is integral to permissive or repressive chromatin conformation, and regulation of gene expression^20^. Expression of *Dot1/Dot1L* is conserved across species^2,6,21–23^ and enrichment of H3K79 methyl marks is associated with actively transcribed chromatin regions^2,6,21–23^. However, DOT1L has also been associated with repression of gene transcription^24^. Thus far, the precise molecular mechanisms of DOT1L regulation of gene expression in HPCs remain undetermined.

We have detected a large number of DEGs in both *Dot1L*-KO or *Dot1L*-MM ESRE cells. Although ~60% of DEGs were common to both *Dot1L*-mutant groups, the remaining ~40% DEGs were unique to either mutant group (**Fig. 3**), which suggest that DOT1L regulates HPCs genes via methyltransferase-dependent and -independent ways. Remarkably, >82% of DEGs in either *Dot1L*-KO or *Dot1L*-MM ESRE cells were found to be upregulated, which indicate that DOT1L primarily acts as a transcriptional repressor in hematopoietic progenitors cells.

Gene ontology and IPA analyses indicated that DOT1L regulates genes, which are responsible for cell signaling and protein-protein interaction. Our previous studies have demonstrated that either loss of DOT1L expression^1,19^ or loss of its methyltransferase activity^3^ leads to G_0_/G_1_ cell cycle arrest and increased apoptosis. IPA analyses showed that the DEGs are linked proliferation and differentiation of HPCs, which were further validated by RT-qPCR analyses. We observed that while genes involved in proliferation of HPCs were significantly downregulated, those involved in induction of differentiation were markedly upregulated (**Fig. 6**). Marked increase in CDK inhibitors positively correlates with the accumulation of cells in G_0_/G_1_ phase and increased proportion of the *Dot1L*-mutant ESRE cells undergoing apoptosis. The gene expression profile also shows a positive correlation with mechanisms involved in incraesed DNA damage.

The most intriguing result of these RNA-seq analyses is the upregulation of >82% genes in *Dot1L*-KO and the *Dot1L*-MM ESRE cells, which proves that DOT1L acts as a transcriptional repressor in HPCs. The other striking result of these analyses is the apparent differences between the *Dot1L*-KO and the *Dot1L*-MM transcriptome profile, suggesting that DOT1L can act in both methyltransferase-dependent and -independent ways.

## Acknowledgements

This work was supported by the National Institutes of Health (grant R01DK091277). The mouse model was generated in the Transgenic and Gene Targeting Institutional Facility of the University of Kansas Medical Center, supported in part by the Center of Biomedical Research Excellence (COBRE) Program Project in Molecular Regulation of Cell Development and Differentiation (NIH P30 GM122731) and the University of Kansas Cancer Center (NIH P30 CA168524).

