## Supplemental Fig. 1 for "DOT1L primarily acts as a transcriptional repressor in hematopoietic progenitor cells"

### Supplemental Figure 1

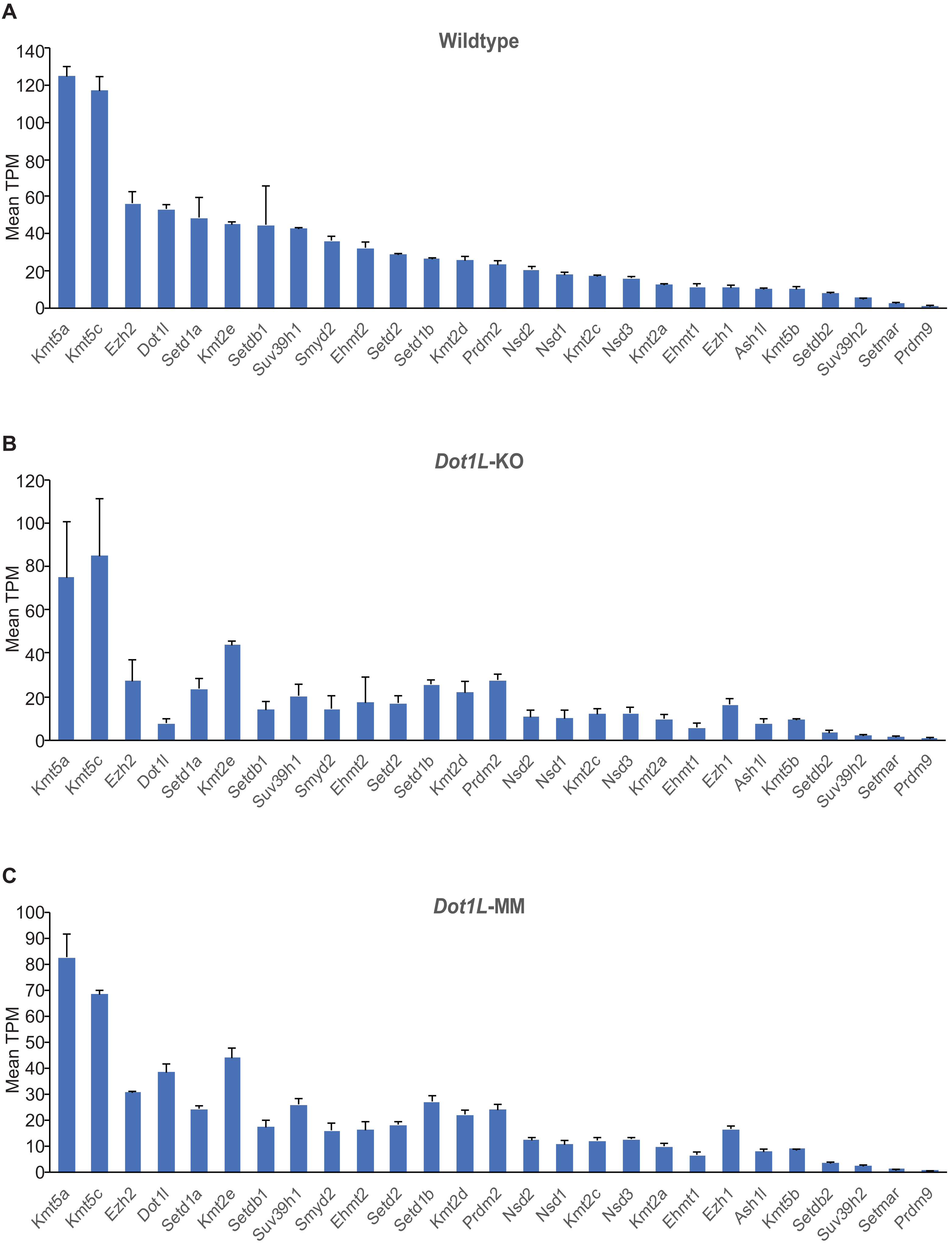

### SUPPL. FIGURE LEGEND

**Suppl. Fig. 1. KMT and KDM profiling analyses in *Dot1L*-KO and *Dot1L*-MM ESRE cells.** The expression levels of a total of 27 KMTs and KDMs were analyzed using our RNA-seq data. The expression levels of *Dot1L* was at the 4$^{th}$ highest level in wildtype (WT) ESRE cells (**A**), whereas it was very low in *Dot1L*-KO cells (**B**) and high in *Dot1L*-MM cells (**C**), comparable to that of WT HPCs.
